# Evaluation of Treatment of Methicillin-Resistant *Staphylococcus aureus* Biofilms with Intermittent Electrochemically-Generated H_2_O_2_ or HOCl

**DOI:** 10.1101/2024.03.22.586337

**Authors:** Md Monzurul Islam Anoy, Won-Jun Kim, Suzanne Gelston, Derek Fleming, Robin Patel, Haluk Beyenal

## Abstract

Chronic wound infections can be difficult to treat and may lead to impaired healing and worsened patient outcomes. Novel treatment strategies are needed. This study evaluated effects of intermittently produced H_2_O_2_ and HOCl, generated via an electrochemical bandage (e-bandage), against methicillin-resistant *Staphylococcus aureus* biofilms in an agar membrane biofilm model. By changing the working electrode potential, the e-bandage generated either HOCl (1.5 V_Ag/AgCl_) or H_2_O_2_ (−0.6 V_Ag/AgCl_). The degree of biocidal activity of intermittent treatment with HOCl and H_2_O_2_ correlated with HOCl treatment time; HOCl treatment durations of 0, 1.5, 3, 4.5, and 6 hours (with the rest of the 6 hour total treatment time devoted to H_2_O_2_ generation) resulted in mean biofilm reductions of 1.36±0.2, 2.22±0.16, 3.46±0.38, 4.63±0.74 and 7.66±0.5 log CFU/cm^2^, respectively vs. non-polarized controls, respectively. However, application of H_2_O_2_ immediately after HOCl treatment was detrimental to biofilm removal. For example, 3-hours HOCl treatment followed by 3-hours H_2_O_2_ resulted in a 1.90±0.84 log CFU/cm^2^ lower mean biofilm reduction than 3-hours HOCl treatment followed by 3-hours non-polarization. HOCl generated over 3-hours exhibited biocidal activity for at least 7.5-hours after e-bandage operation ceased; 3-hours of HOCl generation followed by 7.5-hours of non-polarization resulted in a biofilm cell reduction of 7.92±0.12 log CFU/cm^2^ vs. non polarized controls. Finally, intermittent treatment with HOCl (i.e., interspersed with periods of e-bandage non-polarization) for various intervals showed similar effects (approximately 6 log CFU/cm^2^ reduction vs. non-polarized control) to continuous treatment with HOCl for 3-hours, followed by 3-hours of non-polarization. These findings suggest that timing and sequencing of HOCl and H_2_O_2_ treatments are crucial for maximizing biofilm control.

## INTRODUCTION

Chronic wounds that do not heal through the normal, orderly sequence of repair have low success rates in healing and incur high treatment costs. Pathogens like *Pseudomonas aeruginosa*, *Enterococcus faecalis*, *Staphylococcus aureus*, *Escherichia coli*, *Klebsiella pneumoniae*, *Proteus mirabilis,* and *Acinetobacter baumannii* are commonly found in chronic wounds (1, 2). These microorganism are capable of forming biofilms in which they are encased in a self-secreted extracellular polymeric substance made of polysaccharides, proteins, nucleic acids and lipids (3). Biofilms provide a protective environment for pathogens, enhancing their survival within the wound. This protective mechanism also contributes to delayed wound healing (4–6). It is estimated that 60 to 90% of chronic wounds harbor biofilms (3, 7, 8). Biofilm formation aids microbial survival by restricting the diffusion of molecules and creating an hypoxic environment, which hinders the production of antimicrobial reactive oxygen species by host phagocytes (5, 9, 10). Low oxygen levels in chronic wounds also decrease keratinocyte proliferation (11, 12). Furthermore, biofilms impair epithelialization and granulation tissue formation, leading to delayed wound healing (6). Therefore, targeting biofilms can facilitate the wound healing processes.

Targeting biofilms in wounds is complicated. Biofilms adhere to host tissues, and residing pathogens show enhanced resistance to biocides and antibiotics alike, making higher concentrations necessary for inhibition and eradication than with planktonic forms (13–15). Utilizing a combination of antibiofilm strategies can be beneficial. Shen et al. showed that mechanical agitation, such as ultrasonic or sonic agitation, enhanced the antimicrobial effect of chlorhexidine against biofilm bacteria (16). Wang et al. developed a noninvasive method to eliminate biofilms on metal implants with alternating magnetic fields and ciprofloxacin (17). Tré-Hardy et al. demonstrated that the combination of clarithromycin and tobramycin exhibited synergistic activity against *P. aeruginosa* (18). As such, the possibility of combining biocides is a potential anti-biofilm approach.

The use of hypochlorous acid (HOCl) and hydrogen peroxide (H_2_O_2_) as antimicrobials has been implemented in wound care. Both biocides are found in wound beds; phagocytic cells, including neutrophils and macrophages, undergo oxidative bursts where H_2_O_2_ is generated as part of the inflammatory response (19, 20). Furthermore, approximately 70% of H_2_O_2_ gets converted to HOCl by myeloperoxidase (MPO) enzymes (19, 21). HOCl serves as an innate immune factor and combats infections and foreign biological substances in wound beds. Residual H_2_O_2_ remains active as an antimicrobial, and acts as a signaling molecule and wound healing promoter (22, 23). Thus, HOCl and H_2_O_2_ exhibit favorable antibiofilm and wound healing properties.

Previously, our group developed H_2_O_2_ and (separately) HOCl-generating electrochemical bandages (e-bandages) consisting of carbon fabric working and counter electrodes (CEs) and a silver-silver chloride (Ag/AgCl) pseudo reference electrode to treat biofilms (24, 25). The electrodes were saturated with a phosphate buffered saline (PBS) based hydrogel and separated by layers of cotton fabric. e-Bandage working electrodes (WEs) generate H_2_O_2_ or HOCl based on the applied potential. Briefly, H_2_O_2_ is generated electrochemically on the WE surface of the e-bandage by partial reduction of oxygen at -0.6 V_Ag/AgCl_ (Equation 1) (26). HOCl is generated at 1.5 V_Ag/AgCl_ polarization by oxidation of chloride ions (Equations 2 and 3) (27).

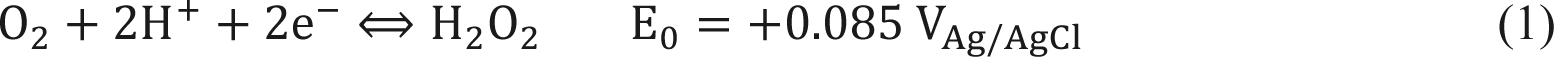

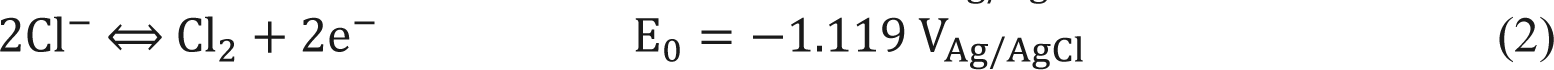

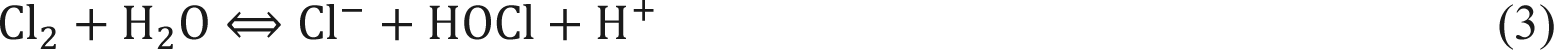

Individually, either showed anti-biofilm effects in an *in vitro* agar membrane biofilm model, *ex vivo* porcine explant biofilm model, and *in vivo* murine wound infection (25, 28, 29). HOCl targets biofilms by decreasing ATP production, and inhibiting bacterial growth, cell division, and protein synthesis, and depressing DNA synthesis (30–32). HOCl deactivates antioxidant enzymes, including catalase, superoxide dismutase, and glutathione peroxidase (28, 33). These antioxidant enzymes act as defense mechanisms for bacteria, allowing them to survive oxidative stress, especially from H_2_O_2_ (34, 35). Although H_2_O_2_ is susceptible to antioxidants, it has the advantage of promoting keratinocyte differentiation and fibroblast migration, augmenting wound healing (36, 37). *In vivo* treatment with H_2_O_2_-producing e-bandages showed that H_2_O_2_ promoted wound healing, as evidenced by decreasing purulence and more rapid wound closure (29). Under similar conditions, HOCl did not improve wound healing, but it did not hinder it (38). It could be beneficial to use both HOCl and H_2_O_2_-generation to treat wound biofilms, with HOCl being used primarily to reduce biofilm cells along with inactivating antioxidant enzymes, and H_2_O_2_ being leveraged to promote wound healing while inhibiting regrowth of biofilms. Such a treatment approach would take advantage of the unique traits of each biocide. However, HOCl reacts with H_2_O_2_ to produce oxygen and water, which might nullify H_2_O_2_ oxidizing effect (28, 39). Therefore, HOCl and H_2_O_2_ should ideally be separated in distance or time. The latter is more suited to the electrochemical approach, as intermittent voltage pulsing is well-established (40). Thus, it would be logical to first produce HOCl for both inactivation of antioxidant enzymes and biocidal effect, and then switch to H_2_O_2_ production for continued biocidal activity and promotion of wound healing.

The goal of this study was to evaluate an intermittent treatment approach employing HOCl and H_2_O_2,_ electrochemically generated using an e-bandage, against a clinical isolate of methicillin-resistant *S. aureus* (MRSA) in an *in vitro* agar membrane biofilm model. It was hypothesized that sequential production of HOCl followed by H_2_O_2_ would result in ideal biofilm targeting. To test this hypothesis, an e-bandage tuned to produce either HOCl or H_2_O_2_ by varying the polarization potential was used. First, the effect of using HOCl for an initial antimicrobial period, and then H_2_O_2_ for the remaining treatment time, was assessed. To assess the impact of intermittent treatment with HOCl- and H_2_O_2_-producing e-bandages, tests were conducted using five time schemes, gradually increasing the HOCl production period. Next, the effect of generation of H_2_O_2_ immediately after HOCl generation was evaluated. To assess this, non-polarized (no production of either agent) control conditions were used by discontinuing e-bandage operation after HOCl generation, with results compared to a system in which H_2_O_2_ was immediately generated after HOCl production. The length of time that electrochemically generated HOCl retained activity after the cessation of e-bandage operation was also determined. Finally, the activity of HOCl intermittently generated for varying time intervals, interspersed with periods of non-polarization, was assessed. Biofilm reduction in terms of colony forming units (CFU) was used to evaluate e-bandage treatments.

## MATERIALS AND METHODS

### Chemical, supplies and bacteria

Potassium chloride (KCl) (cat#3040-01), and potassium phosphate monobasic (KH_2_PO_4_) (cat#3246-05) were purchased from J.T. Baker. Sodium phosphate dibasic dehydrate (HNa_2_O_4_P·2H_2_O) (cat#101835432) and sodium hypochlorite solution (NaOCl) (CAS #7681-52-9) were purchased from Sigma-Aldrich. Sodium chloride (NaCl) (cat#S271-3), tryptic soy broth (TSB) (cat#DF0370-13-3), and tryptic soy agar (TSA) (cat#DF0369-17-6) were purchased from Fisher Scientific. MRSA IDRL-6169, a clinical prosthetic hip isolate that is resistant to methicillin and mupirocin, was used for all experiments.

### e-Bandage

Construction and preparation of the e-bandages was as previously described (28). e-Bandages were made of three electrodes: 1.77 cm^2^ of carbon fabric (Panex 30 PW06, Zoltek Companies, Inc.) as the WE and CE, and an Ag/AgCl wire as the pseudo reference electrode. The electrodes are separated by cotton fabric and held together using silicon adhesive. 30 AWG titanium wires (TEMco, Amazon.com, cat#RW0517) were pressed with nylon sew-on snaps (Dritz, Spartanburg, SC, item#85) for electrical connectivity between the potentiostat cable (Interface 1010T/Interface 1000, Gamry) with a multiplexer (IMX8/ECM8, Gamry) and the WE and CE. The CE and WE were covered with hydrogel made from 1x PBS mixed with 1.8 % (w/v) xanthan gum (Namaste Foods, Amazon.com, UPC: 301155217160) to ensure electrochemical connectivity.

### *In vitro* agar membrane biofilm model

For the *in vitro* agar membrane biofilm model, a generation 1 streak plate was prepared by streaking frozen stock culture on a tryptic soy agar (TSA) plate. The generation 1 plate was incubated at 37°C for 24 hours. A generation 2 plate was made by streaking a colony picked from the generation 1 plate on a fresh TSA plate. The generation 2 plate was incubated at 37°C for 24 hours. A bacterial broth was made by adding a single colony from the generation 2 plate to a 2 mL TSB solution. The broth culture was placed in the incubator at 37°C at 150 rpm until it reached a turbidity of 0.5 McFarland (approximately 2-4 hours). Finally, 2.5 µL of this broth was spotted on the center of a UV-sterilized 13 mm polycarbonate membrane (Whatman Nuclepore polycarbonate hydrophilic membranes, Cytiva #10417001) atop a TSA plate. Biofilms were grown on the membranes for 24 hours at 37°C.

### e-Bandage treatment

e-Bandage treatment was done with the agar membrane biofilm model. After 24 hours of growth, biofilms were transferred to a fresh TSA plate. One hundred microliters of hydrogel were placed on top of biofilm, and an e-bandage was placed on top of the hydrogel, with the WE side in contact with the biofilm. One hundred microliters of hydrogel were placed in between the cotton fabric layers of the e-bandage, where the reference electrode was situated, with a syringe needle. Finally, another 100 µl of hydrogel was placed on top of the e-bandage. A sterile Tegaderm™ transparent film (3 M, ref #1622 W) was used to cover the e-bandage. e-Bandage wires were taped to the side of the TSA plate. Finally, the lid of the TSA plate was placed, and the TSA plates wrapped with parafilm. Each e-bandage was connected to a potentiostat (Interface 1010T/Interface 1000, Gamry) with a multiplexer (IMX8/ECM8, Gamry). The WE was polarized to -0.6 V_Ag/AgCl_ or 1.5 V_Ag/AgCl_ to generate H_2_O_2_ or HOCl, respectively, on the WE surface. For non-polarized (no production of oxidizing agents) conditions, the potentiostat was set to idle mode. **Fig. 1A** shows a schematic of the *in vitro* experimental setup with an e-bandage placed on top of a laboratory-grown biofilm on a polycarbonate membrane, and the potentiostat conditions for generation of HOCl or H_2_O_2_, or the non-polarized state.

**Figure 1.**
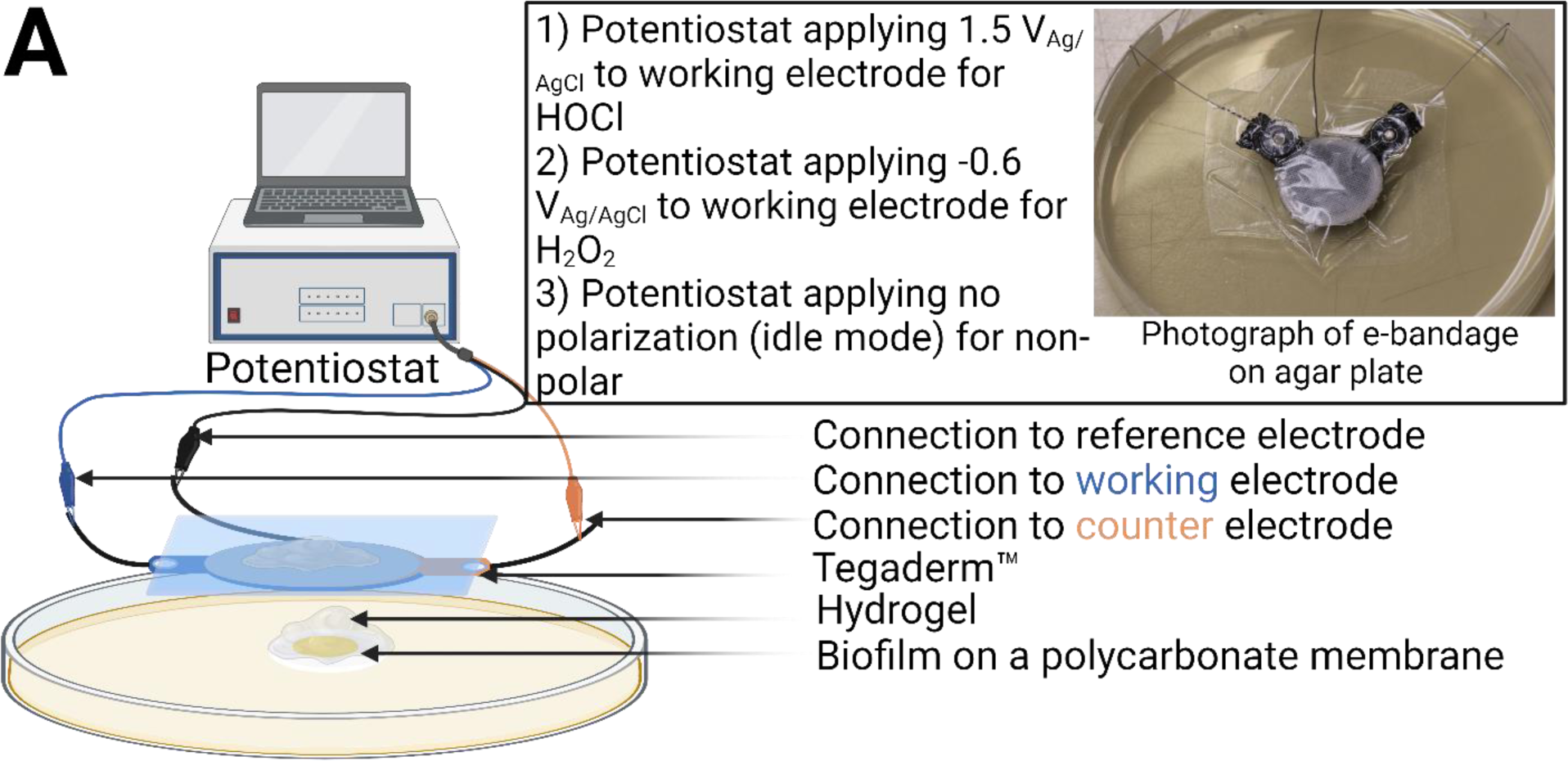
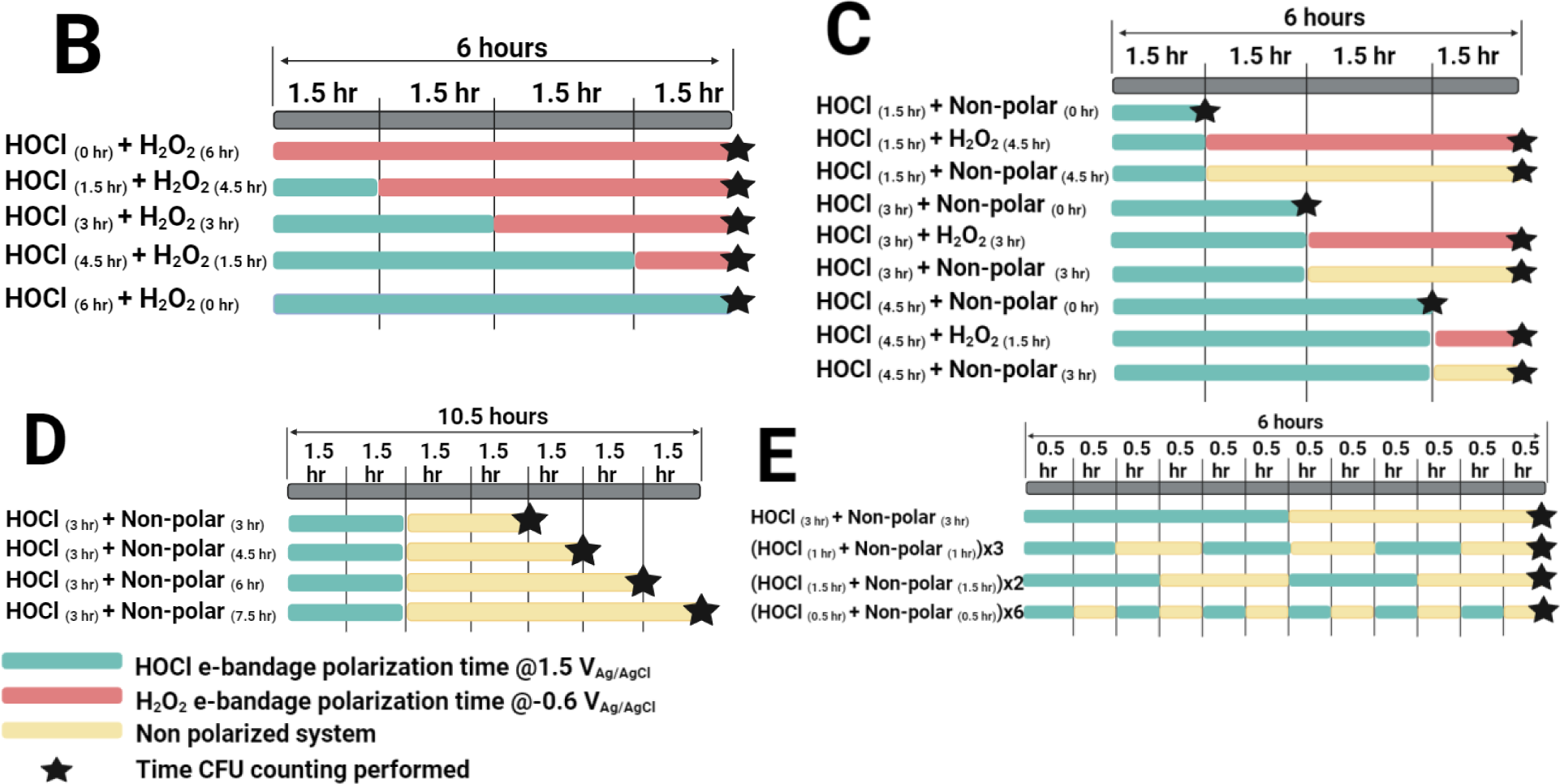
Schematic of *in vitro* experimental setups and treatment conditions. **A** *in vitro* experimental setup with e-bandages connected to a potentiostat, placed on top of laboratory grown MRSA IDRL-6169 biofilms on a TSA plate. Inset) Photograph of an e-bandage on agar plate. Figures are not drawn to scale. B-E show treatment length and type, waiting periods, and times at which CFU counting was performed. **B)** Representation of conditions for intermittent treatment with HOCl and H_2_O_2_; treatments investigated effects of varying HOCl and H_2_O_2_ treatment times. **C)** Representation of the experimental conditions for testing effects of intermittent HOCl and H_2_O_2_ exposure compared to HOCl exposure with periods of non-polarization. Treatments examined whether immediate treatment with H_2_O_2_ following HOCl treatment is beneficial for biofilm control. **D)** Representation of experimental conditions for testing the duration of effect of HOCl e-bandage followed by non-polarization. Treatments explored the duration for which HOCl exhibits antimicrobial activity within the system following discontinuation of e-bandage polarization. **E)** Representation of experimental conditions for testing the effect of intermittent production of HOCl. Treatments compared the effectiveness of intermittent HOCl treatment for different time intervals.

### Experimental setup

An *in vitro* experimental setup was utilized to investigate the impact of various treatment protocols with electrochemically generated HOCl or H_2_O_2_ using an e-bandage against MRSA agar membrane biofilms (**Fig. 1A**). Intermittent treatment with HOCl and H_2_O_2_, was tested using five treatment protocols with 6 hours of total treatment time each (**Fig. 1B**). These protocols included 6 hours of H_2_O_2_, 6 hours of HOCl, 3 hours of HOCl followed by 3 hours of H_2_O_2_, 1.5 hours of HOCl followed by 4.5 hours of H_2_O_2_, and 4.5 hours of HOCl followed by 1.5 hours of H_2_O_2_ production. H_2_O_2-_production was compared to non-polar conditions (**Fig. 1C**). Duration of the biocidal effect of e-bandage HOCl production followed by periods of non-polarization of varying durations was also tested (**Fig. 1D**). Specifically, three-hours of HOCl production was followed by 3, 4.5, 6, and 7.5 hours of non-polarization. To test effects of intermittent HOCl production alone, cycles of HOCl and non-polarization were tested at various intervals and compared to a 6-hour non-polarized control group (**Fig. 1E**).

### Colony forming unit (CFU) quantification after e-bandage treatment

Biofilms (initial, control, and e-bandage treated) were quantified by calculating CFUs as described previously (24, 28, 29). Briefly, e-bandages were removed from TSA plates and placed in Petri dishes. Biofilms were scraped from the WE surface into five mL of PBS. The PBS solution and the membranes were transferred to 15 mL centrifuge tubes, vortexed for 2 min, and sonicated for 10 min. Cells were then pelleted via centrifugation at 2910 relative centrifugal force (rcf) for 10 minutes and resuspended in 1 mL of PBS. The suspension was serially diluted (10-fold) and 10 μl of each dilution spotted on TSA and incubated at 37°C overnight. Colonies were counted after 24 hours.

### HOCl concentration profiles

HOCl microelectrodes were constructed by following a published protocol (41). Briefly, a two-electrode system, including a platinum working electrode to sense HOCl concentration, and an external reference electrode (Ag/AgCl in saturated KCl) were used. To construct the HOCl-sensing microelectrode, a 50 µm platinum wire was electrochemically etched and inserted into a glass capillary (Corning 816). The glass capillary was sealed, and the tip was exposed. A 20-30 µm diameter platinum ball was electrodeposited at the microelectrode tip. The platinum ball at the tip of the microelectrode was coated with 5% cellulose acetate membrane and dried for 24 hours. The microelectrode was polarized at 0.35 V_Ag/AgCl_ using a Gamry 1010E potentiostat (Gamry Instruments). Calibration was done using a NaOCl standard solution in PBS before and after each concentration profile measurement. The microelectrode had limit of 11µM (S/N=2) and a response time of less than three seconds.

An e-bandage was soaked in PBS and 100 µL of hydrogel was applied between the cotton fabric layers. The e-bandage was placed upside down (the WE facing up) onto a TSA plate. Although this is not how e-bandage operates in practice, this setup allowed use of a microelectrode for measurements on the WE. However, this also limited measurement time since hydrogel was open to the air, leading to gradual evaporation of the water content. It was found that microelectrodes could be operated for up to two hours without being affected by evaporation. One hundred µL of hydrogel was applied to the WE. The microelectrode tip was placed 100 µm above the WE surface within the hydrogel using a stepper motor controlled by custom LabVIEW software (Physik Instruments, PI M-230.10 S). The WE was polarized using a benchtop potentiostat (G300, Gamry Instruments) to generate HOCl or H_2_O_2_. HOCl was generated by polarizing the WE at 1.5 V_Ag/AgCl_, and H_2_O_2_ was generated by polarizing the WE at -0.6 V_Ag/AgCl_. Two conditions were tested to verify the change in HOCl concentration when the WE polarization was switched from 1.5 V_Ag/AgCl_ to -0.6 V_Ag/AgCl_ (from HOCl generation to H_2_O_2_ generation); 1) 1 hour of HOCl followed by 1 hour of H_2_O_2_ produced by the WE, and 2) 1 hour of HOCl polarization followed by 1 hour of non-polarization (e-bandage is inactive).

### Statistics

Data were displayed as individual data points for at least four biological replicates. A two-sided Wilcoxon rank-sum test was performed to compare log reduction of bacterial counts associated with e-bandage treatment to untreated controls, unless otherwise indicated. Statistical analysis and figures were generated using MATLAB® (version R2021b).

## RESULTS AND DISCUSSION

### Intermittent treatment with HOCl and H_2_O_2_ generated by an e-bandage shows increasing biocidal efficacy with longer HOCl treatment times

Intermittent treatment conditions using HOCl and H_2_O_2_ e-bandage were tested against MRSA biofilms. Six hours of treatment with HOCl-producing e-bandage reduced the initial bacterial load to 2.21 ± 0.5 log CFU/cm^2^ (7.45 ± 0.52 log CFU/cm^2^ reduction; **Supplementary Fig. S1**). Six hours of treatment using an H_2_O_2_-producing e-bandage showed only 1.41 ± 0.22 log CFU/cm^2^ reduction. Four hours of treatment with individually generated HOCl and H_2_O_2_ was also tested, with both HOCl and H_2_O_2_ showing less efficacy than 6 hours of treatment with either alone. Given that six hours of treatment with an HOCl-generating e-bandage showed maximum CFU reduction against MRSA while still falling above the limit of detection (LOD), this treatment time was used for subsequent intermittent therapy experiments. Initial biofilm loads averaged 9.69 ± 0.1 log CFU/cm^2^, and non-polarized controls averaged 9.87 ± 0.03 log CFU/cm^2^ after 6 hours (Fig. 2). Increasing the duration of HOCl treatment showed time-dependent increases in biofilm reduction. When MRSA biofilm was treated with an HOCl-generating e-bandage for 0, 1.5, 3, 4 and 6 hours of the total 6 hour treatment time (i.e., with the rest of the treatment being with an H_2_O_2_-generating e-bandage), averages of 1.36 ± 0.2, 2.22 ± 0.17, 3.46 ± 0.38, 4.63 ± 0.74 and 7.66 ± 0.5 log CFU/cm^2^ reduction, respectively, were observed. Thus, intermittent treatment with HOCl and H_2_O_2_-producing e-bandages against MRSA biofilms showed an increase in biocidal efficacy with longer periods of HOCl treatment. Previously, we reported that treatment with HOCl producing e-bandages showed a time dependent response against MRSA biofilms, with no viable cells detected in broth culture after 12 hours of treatment (24). In contrast, 48 hours of treatment with an H_2_O_2_-producing e-bandage was necessary to eradicate MRSA biofilms (25). This is supported by results of the tested intermittent treatment conditions against MRSA biofilms reported here. Results of this study inform selection of a minimal treatment time, at which anti-biofilm activity is demonstrated, without risking harm to host tissue through prolonged exposure to HOCl (42).

**Figure 2.**
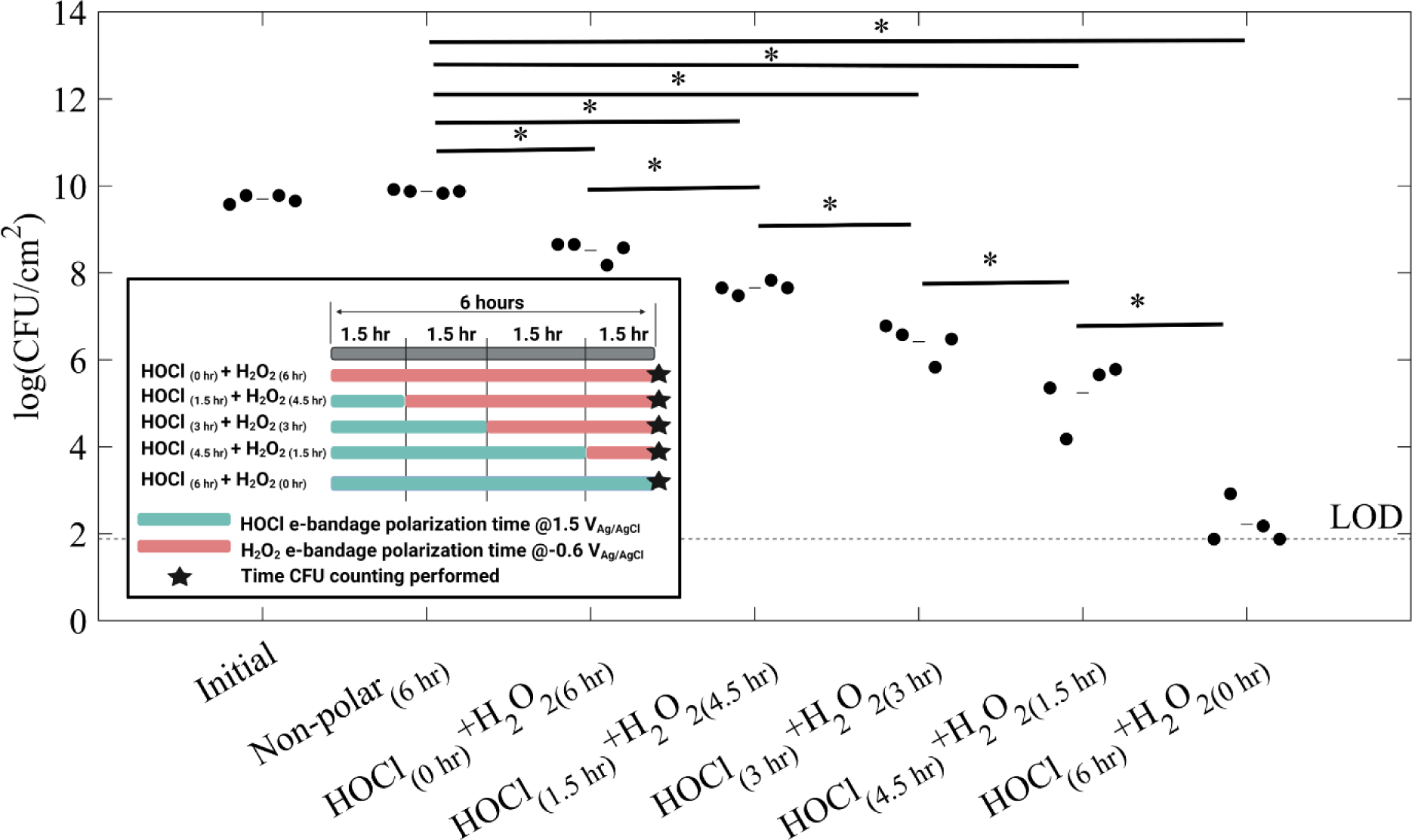
Effects of intermittent treatment with HOCl and H_2_O_2_ produced by an e-bandage against MRSA biofilms. Five treatment conditions were tested. “Non-polar” refers to an inactive e-bandage. Subscripts show the duration of the treatment. Biofilm cell counts are shown as log CFU/cm^2^, with treatments compared with untreated controls (non-polar). Additionally, each treatment condition was compared with adjacent treatment conditions. Data is represented as individual data points (circles) and means (lines) of at least four independent biological replicates (n ≤ 4, *P <0.05, two-sided Wilcoxon rank-sum test). LOD = limit of detection. Inset) Schematic showing treatment length and type, waiting periods, and times CFU counting was performed. Overall, intermittent treatment with HOCl and H_2_O_2_ generated via an e-bandage showed increasing biocidal efficacy with longer HOCl treatment duration.

### H_2_O_2_ treatment immediately after HOCl treatment mitigates HOCl effects

The effect of H_2_O_2_ treatment after HOCl treatment was evaluated by polarizing the e-bandage at 1.5 V_Ag/AgCl_ (for generating HOCl) for 1.5, 3 and 4.5 hours, followed by -0.6 V_Ag/AgCl_ (for generating H_2_O_2_) for the remaining time (total treatment time, 6 hours). For control groups, the e-bandage was non-polarized instead of polarizing for H_2_O_2,_ following HOCl production for the remaining time of each treatment. 4.5 hours of HOCl treatment with 1.5 hours of non-polarization was then compared to 4.5 hours of HOCl treatment followed by 1.5 hours of H_2_O_2_ treatment to test for potential detrimental effects of H_2_O_2_ on HOCl efficacy. Fig. 3A shows that after polarizing for 1.5 hours at 1.5 V_Ag/AgCl_ for HOCl treatment, an average of 7.14 ± 0.68 log CFU/cm^2^ of MRSA was observed. Similarly, after 1.5 hours of HOCl treatment, followed by non-polarization for 4.5 hours, an average of 7.65 ± 0.014 log CFU/cm^2^ was observed. This non-significant (P=0.4857) increase in bacterial counts indicates that following HOCl generation over 1.5 hours, the system maintains a fixed amount of bacteria for at least 4.5 hours. Conversely, an average of 8.5 ± 0.21 log CFU/cm^2^ was observed after treating biofilms with 1.5 hours of HOCl followed by 4.5 hours of H_2_O_2_. This significant increase (P=0.0286) in viable cell counts compared to treatment with 1.5 hours of HOCl alone indicates that generation of H_2_O_2_ for 4.5 hours after 1.5 hours of HOCl generation negates the effects of HOCl production to a degree. This detrimental effect of immediate H_2_O_2_ treatment after HOCl treatment was observed for both 3 and 4.5 hours of HOCl treatment (Fig. 3B and 3C). Specifically, 3 hours of HOCl treatment alone resulted in an average of 5.1 ± 0.4 log CFU/cm^2^ remaining, while adding 3 hours of non-polarization after the 3 hours of HOCl treatment reduced the bacterial count further to 4.52 ± 0.66 log CFU/cm^2^. This significant reduction (P=0.0286) may be due to residual HOCl remaining in the hydrogel. However, three hours of treatment with H_2_O_2_ following 3 hours HOCl treatment yielded a higher bacterial load of 6.4 ± 0.4 log CFU/cm^2^ compared to either HOCl alone or HOCl followed by non-polarization. 4.5 hours of HOCl treatment with and without 1.5 hours of subsequent non-polarization resulted in 4.73 ± 1.07 and 2.65 ± 0.89 log CFU/cm^2^ respectively. Adding 1.5 hours of H_2_O_2_ treatment instead of non-polarization decreased treatment efficacy, resulting in an average of 5.24 ± 0.73 log CFU/cm^2^ remaining.

**Figure 3.**
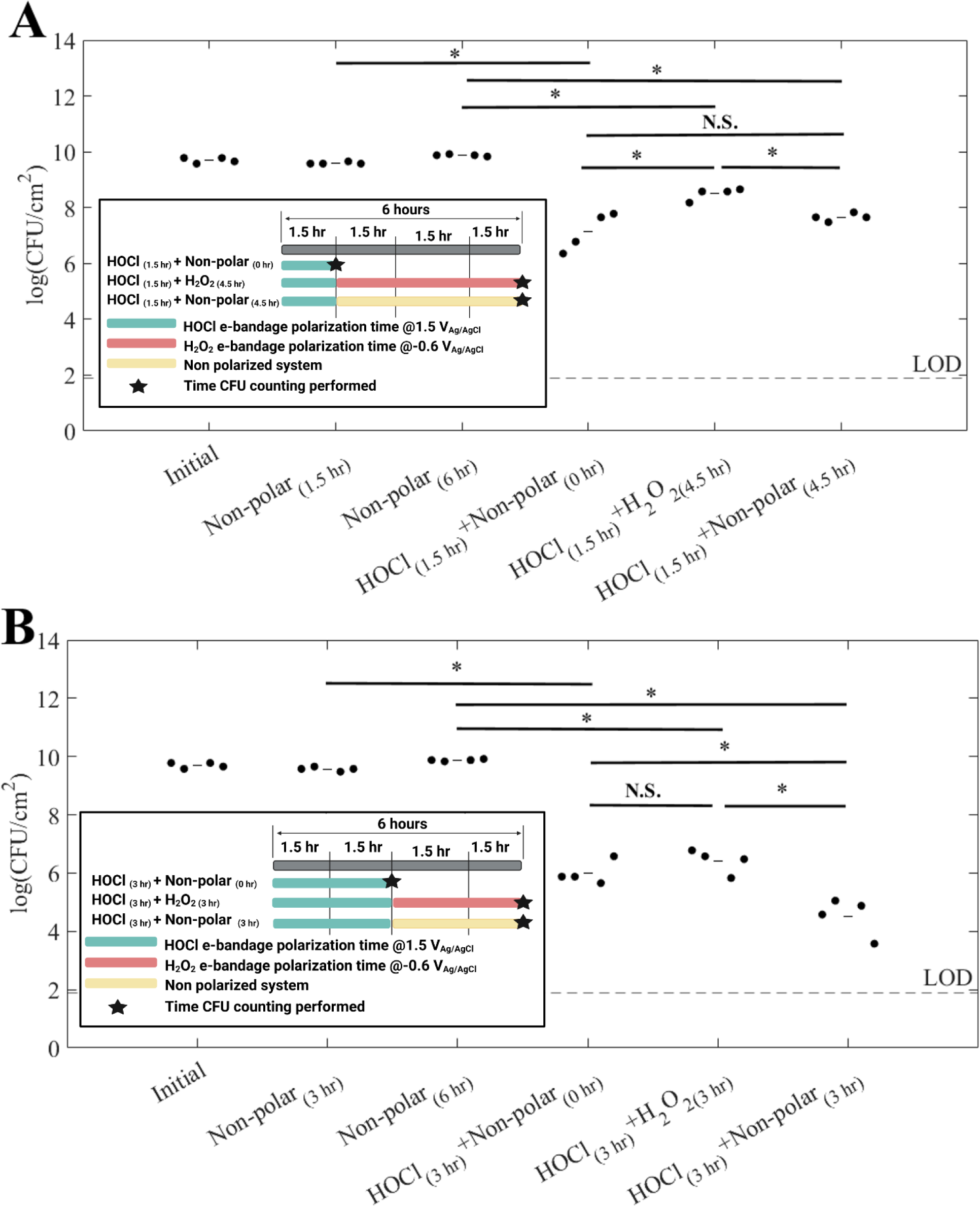
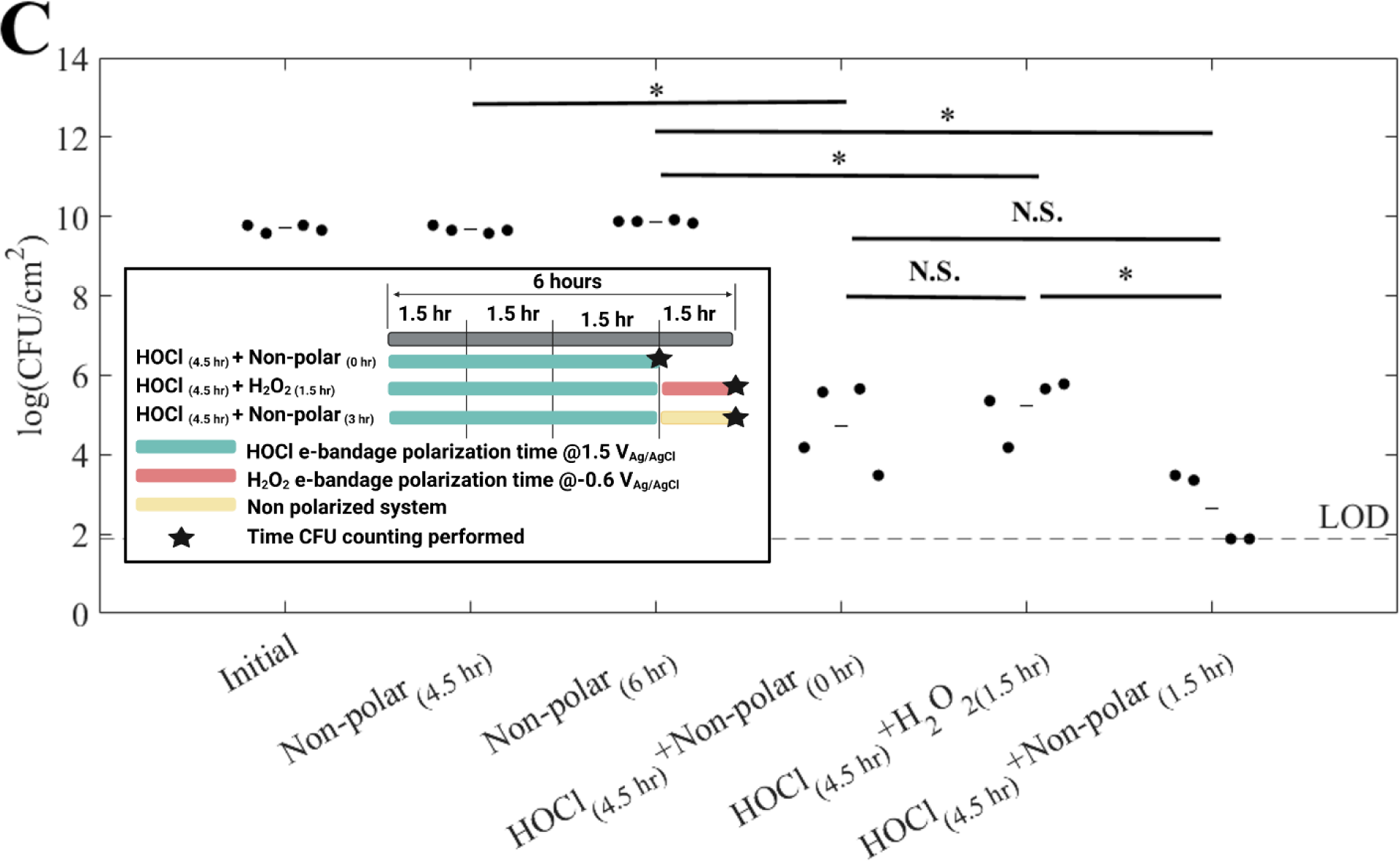
Immediate treatment using H_2_O_2_ after HOCl reduces the efficacy of e-bandages against MRSA biofilm. Treatment using H_2_O_2_ after HOCl with e-bandage was tested at 3 time points depending on the HOCl treatment time: A) 1.5 hours, B) 3 hours and C) 4.5 hours. Total treatment time was 6 hours. “Non-polar” refers to an inactive e-bandage. Subscripts show the duration of the treatment. Biofilm counts are shown as log CFU/cm^2^; treatments were compared with untreated controls. Additionally, each treatment condition was compared with each other. Data is represented as individual data points (circles) and means (lines) of at least four independent biological replicates (n ≤ 4, *P <0.05, N.S. = non-significant, two-sided Wilcoxon rank-sum test). LOD = limit of detection. Inset A, B, & C) Schematic showing treatment length and type, waiting periods, and times CFU counting was performed. Treatment using electrochemically generated H_2_O_2_ immediately after HOCl treatment showed detrimental effects.

These results indicate that treatment using electrochemically generated H_2_O_2_ is not desirable immediately after HOCl generation for biofilm control. Electrochemical generation of these species depends on polarization potential; this electrochemical process is beneficial because it allows for low concentrations to be generated over long periods of time without being cytotoxic (41, 43). Since longer e-bandages treatment times would result in accumulation of higher concentrations of these species, shorter treatment times may be desirable. However, the two oxidant species are reactive, and when contacting one other, react to create singlet oxygen and water (44, 45). Indeed, kinetic studies have shown that HOCl and H_2_O_2_ react as a second order reaction, with a reaction rate of 195.5 ± 3.3 M^-1^s^-1^, with production of single oxygen happening almost immediately (39). Although this oxygen species is favorable to biofilm control, here it appeared to be less effective than HOCl alone (46). Finally, these experiments showed that the hydrogel used may retain the electrochemically generated HOCl and carry the effect over into non-polar conditions for residual bactericidal activity. Because of this, care must be taken to design intermittent treatment protocols that consider the presence of residual HOCl. In the next section, the efficacy of hold time of the electrochemically generated HOCl to improve timing of intermittent treatment was evaluated.

We previously measured and reported HOCl and H_2_O_2_ concentrations near e-bandages WEs, when produced individually (24, 47). When the WE is polarized to 1.5 V_Ag/AgCl_ or -0.6 V_Ag/AgCl_, it generates HOCl (24) or H_2_O_2_, respectively (47). However, it has not been shown what happens to HOCl concentrations when the WE is polarized for H_2_O_2_ generation immediately after HOCl generation. For this purpose, in the first experiment, e-bandage was polarized at 1.5 V_Ag/AgCl_ to generate HOCl at the WE surface for 1 hour, followed immediately by 1 hour of H_2_O_2_ genearation by changing the polarization of the WE from 1.5 V_Ag/AgCl_ to -0.6 V_Ag/AgCl_. Fig. 4 shows that polarization of WE to 1.5 V_Ag/AgCl_ generates HOCl. HOCl concentrations gradually increase, reaching ∼ 140 mM within 40 minutes. Later, the concentration of HOCl decreases due to diffusion and possible reaction with organics in the hydrogel, reaching ∼ 100 mM (48). However, immediately upon polarization of the WE at -0.6 V_Ag/AgCl_ (to generate H_2_O_2_), HOCl concentrations dropped below the detection limit, due to reaction between HOCl and H_2_O_2_ (Equation 4) (28, 44, 45).

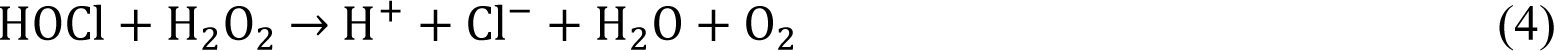

**Figure 4.**
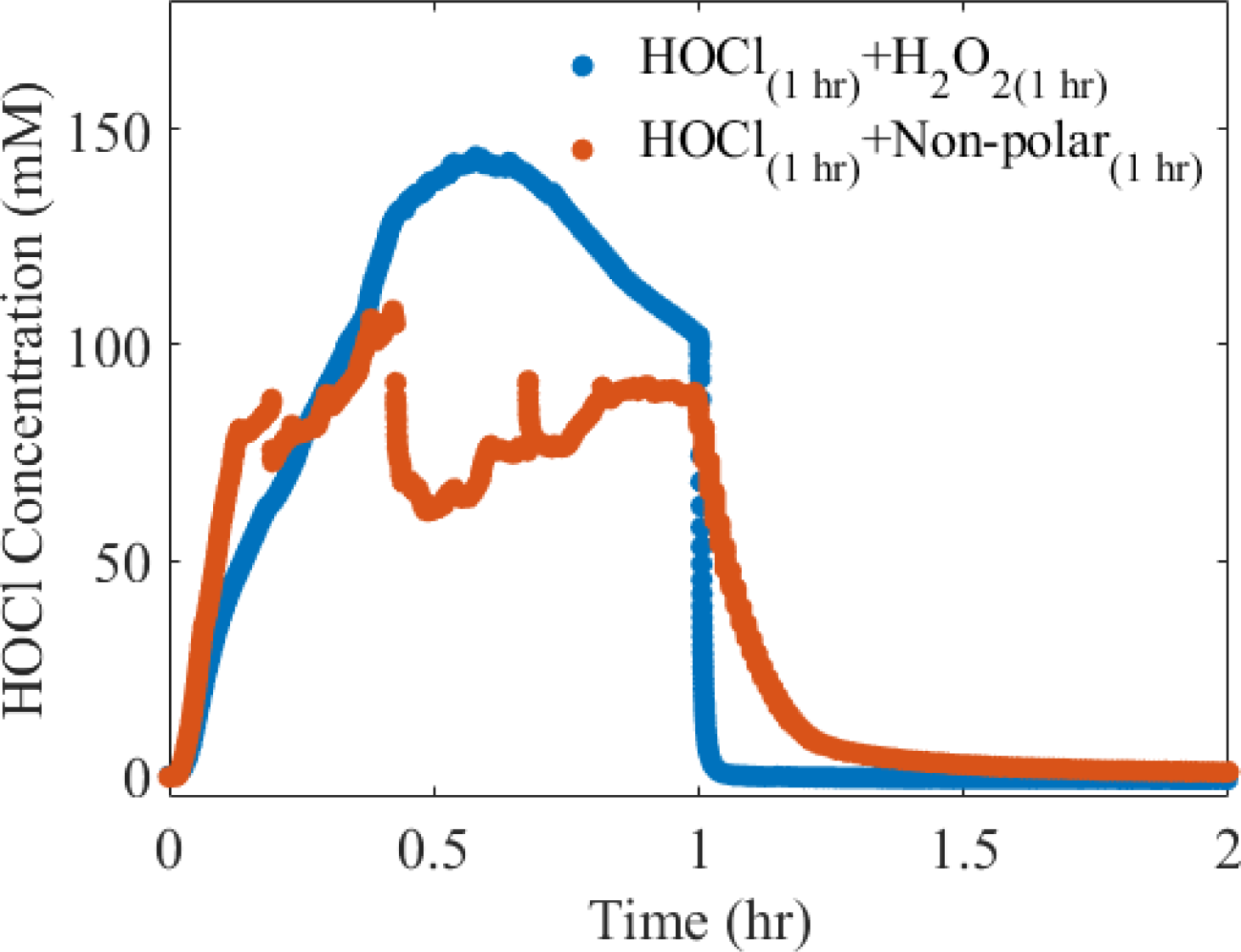
HOCl concentration profiles one hundred µm from the WE surface, as measured by an HOCl microelectrode. “Non-polar” refers to an inactive e-bandage. Subscripts show the duration of the compound generated (HOCl or H_2_O_2_). HOCl_(1 hr)_+H_2_O_2(1 hr)_: the e-bandage was polarized at 1.5 V_Ag/AgCl_ (for generation of HOCl on the WE of e-bandage) for one hour and switched to - 0.6 V_Ag/AgCl_ (for generation of H_2_O_2_ on the WE of e-bandage) for one hour. HOCl_(1 hr)_+Non-polar_(1 hr)_: the e-bandage was polarized at 1.5 V_Ag/AgCl_ for one hour followed by one hour of non-polarization. Generation of H_2_O_2_ immediately after HOCl shows an immediate decrease in HOCl concentration due to the reaction between HOCl and H_2_O_2_.

In a subsequent experiment (Fig. 4), instead of generating H_2_O_2_ after 1 hour of HOCl generation, WE polarization was stopped and the concentration of HOCl monitored. Nealrly 60 minutes elapsed before the HOCl concentration decreased to below limit of detection. The data in Fig. 4 shows that 1) H_2_O_2_ generation after HOCl generation removes generated HOCl, and 2) if HOCl is not generated continuously, it degrades over time. There were differences in maximal concentrations are likely due to difficulty positioning the microelectrode tip at exactly 100 µm from the carbon fabric surface (Fig. 4).

### Treatment with electrochemically generated HOCl retains sustained biofilm reductions after ceasing e-bandage operation

To assess the activity of the e-bandage system after stopping polarization, the e-bandage was polarized at 1.5 V_Ag/AgCl_ to generate HOCl for 3 hours, followed by leaving the e-bandage non-polarized for 3, 4.5, 6 and 7.5 hours. Figure 5 shows that treatment with electrochemically generated HOCl left undisturbed in a non-polarized system for 3, 4.5, 6 and 7.5 hours maintained reduction of biofilms in all cases (P=0.0286) with resulting CFU quantities of 4.52 ± 0.66, 3.56 ± 0.89, 3.21 ± 0.90 and 1.95 ± 0.15 log CFU/cm^2^, respectively. Increased CFU reduction was correlated with increased non-polarization periods, suggesting effects of retained electrochemically generated HOCl (Fig. 5). Overall, 3 hours of electrochemical generation of HOCl followed by non-polarization exhibited a similar antimicrobial effect to 6 hours of HOCl treatment, suggesting that continuous HOCl generation may not be needed.

**Figure 5.**
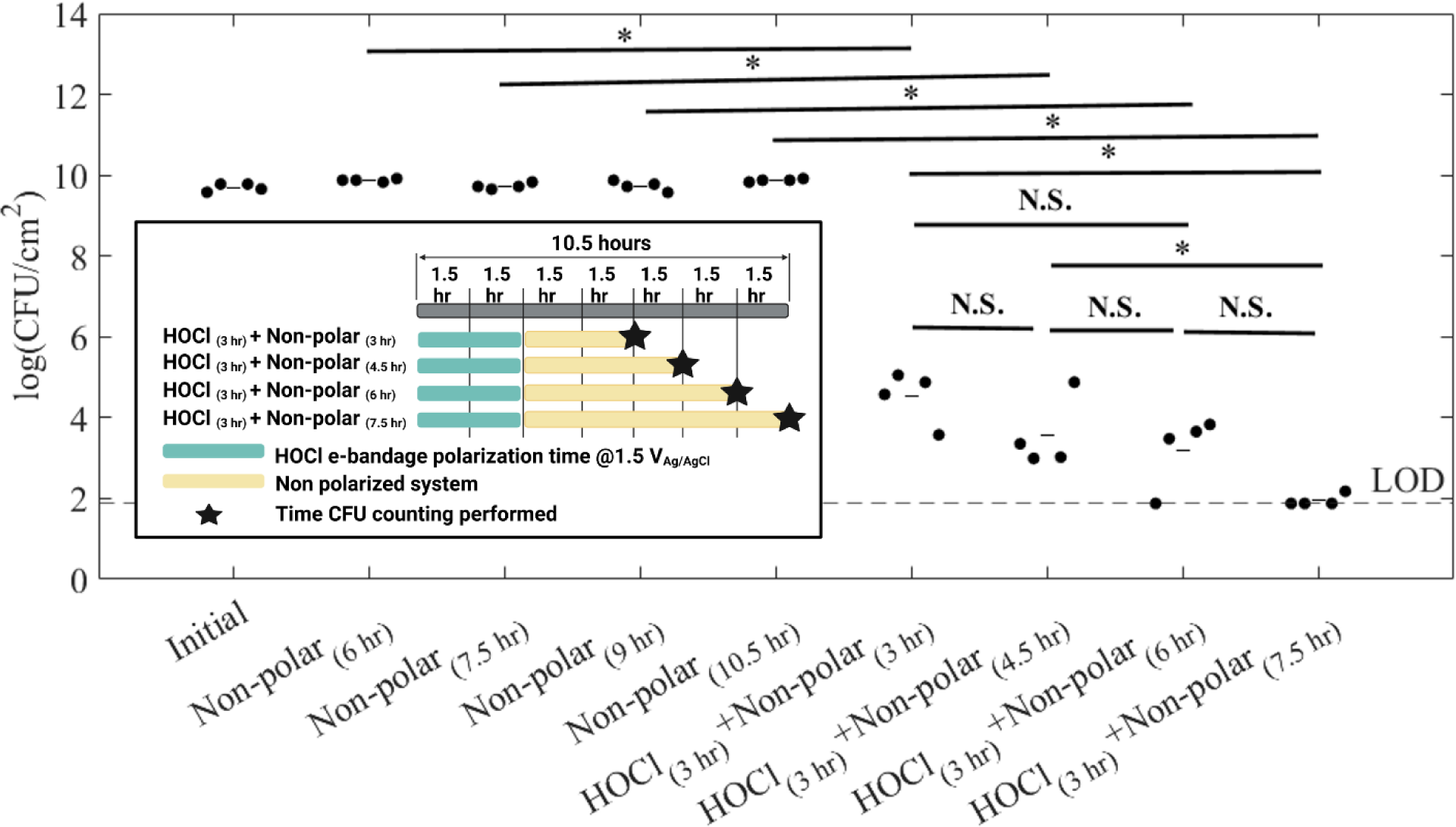
Electrochemically generated HOCl shows sustained biocidal activity in controlling MRSA in an agar membrane biofilm model. “Non-polar” refers to an inactive e-bandage. Subscripts show the duration of the treatment. Biofilm cell counts are shown as log CFU/cm^2^, and treatments are compared with untreated controls (non-polar). Additionally, each treatment condition is compared with each other. Data is represented as individual data points (circles) and means (lines) of at least four independent biological replicates (n ≤ 4, *P <0.05, N.S. = non-significant, two-sided Wilcoxon rank-sum test). LOD = limit of detection. Inset) Schematic showing treatment length and type, waiting periods, and times CFU counting was performed. Three hours of HOCl generation through the e-bandage yielded persistent activity for at least 7.5 hours after generation.

### Intermittent treatment using HOCl, irrespective of time interval, yielded similar effects to continuous treatment

Given that 3 hours of electrochemically generated HOCl showed sustained biofilm reductions after a non-polarized period, intermittent treatment using HOCl delivered at various intervals was tested. Various time intervals were used to evaluate efficacy of intermittent treatment using HOCl-generating e-bandages. Each treatment time was six hours, and each involved three hours total of HOCl generation. Three different protocols were tested: 1) Three rounds of 1-hour HOCl + 1-hour non-polar; 2) Two rounds of 1.5 hours HOCl + 1.5 hours non-polar; and 3) Six rounds of 0.5 hours HOCl + 0.5 hours non-polar. Figure 6 shows that each approach yielded significant biofilm reductions (P=0.0286) compared to untreated controls. Although there were no significant differences between effects of the three protocols, 1.5 hours HOCl + 1.5 hours non-polar × 2 showed the highest average biofilm reduction (3.50 ± 1.88 log CFU/cm^2^).

**Figure 6.**
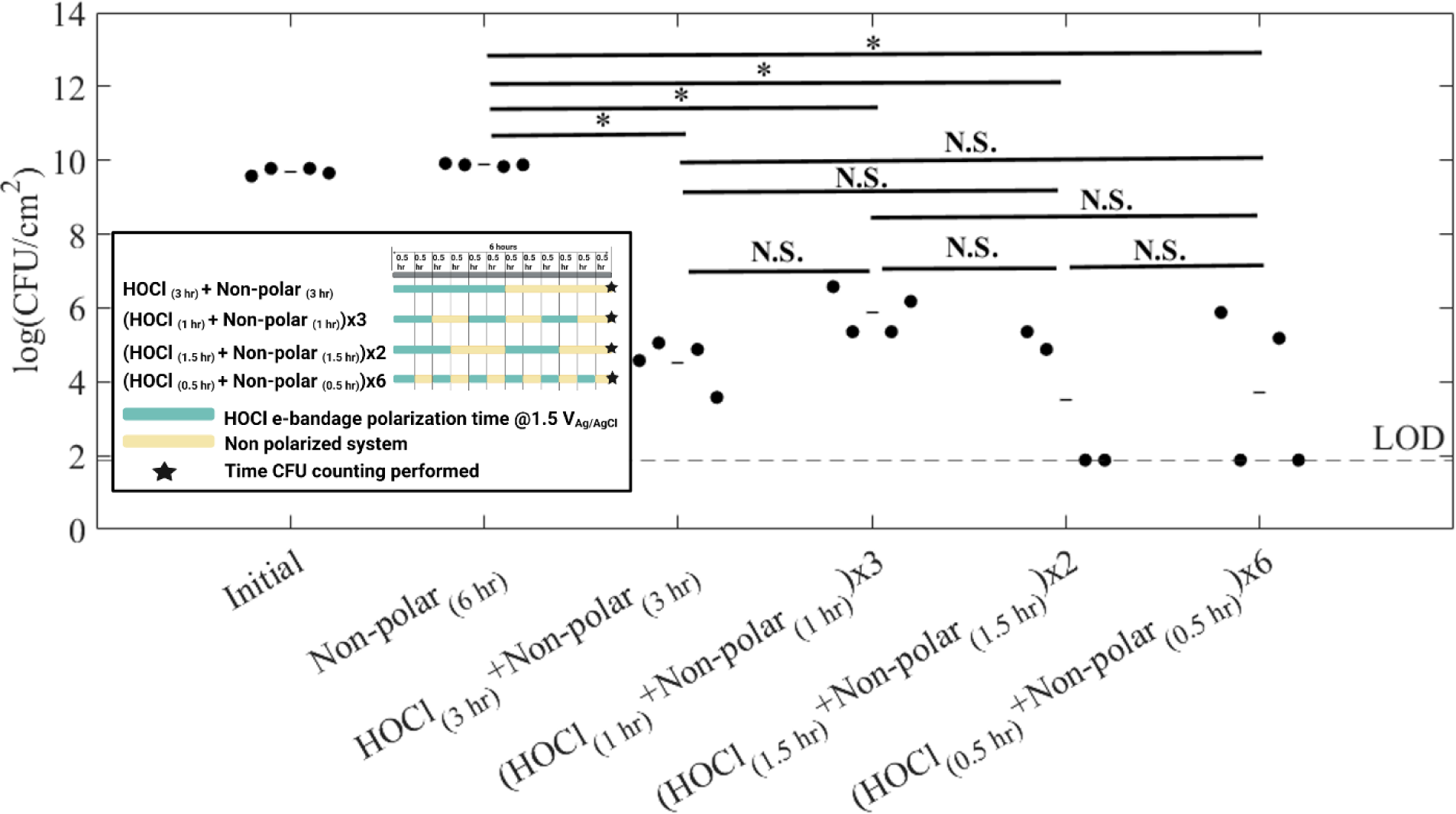
Intermittent HOCl treatment using the e-bandage system shows biocidal activity. “Non-polar” refers to an inactive e-bandage. Subscripts show the duration of the treatment. “x Number” shows the number of cycles. Biofilm cell counts are shown as log CFU/cm^2^, and treatments were compared to untreated controls (non-polar). Additionally, each treatment condition was compared with one another. Data is represented as individual data points (circles) and means (lines) of at least four independent biological replicates (n ≤ 4, *P <0.05, N.S. = non-significant, two-sided Wilcoxon rank-sum test). LOD = limit of detection. Inset) Schematic showing treatment length and type, waiting periods, and times CFU counting was performed. Intermittent treatment using HOCl for half of the treatment period showed similar effects regardless of time intervals.

### Practical Implications

Both HOCl and H_2_O_2_ are antimicrobial agents. The described e-bandage is an innovative potential wound dressing that can deliver either of these compounds *in situ* at the wound site. Given the highly reactive nature of these compounds and their propensity for continuous degradation, electrochemical generation is advantageous in maintaining consistent concentrations (49, 50). Electrochemical generation of HOCl or H_2_O_2_ using e-bandages has demonstrated efficiency in controlling biofilms individually (51, 52). Furthermore, treatment using electrochemically generated H_2_O_2_ or HOCl did not show emergence of resistance for *S. aureus* or *P. aeruginosa* biofilm over 10 iterations (15). However, to effectively utilize both H_2_O_2_ and HOCl-producing e-bandage, a strategic approach is needed. Here, MRSA, a common wound pathogen, was studied (53). Results demonstrate strategies to minimize exposure to HOCl while maximizing antimicrobial efficacy. An important finding was that H_2_O_2_ should not be deployed immediately after HOCl. A separate study has shown that an H_2_O_2_ e-bandage accelerates wound closure and healing in an *in vivo* environment, while also reducing biofilms (29). In light of these findings, a novel treatment regimen can be considered. Initially, treatment using HOCl can be utilized, followed by a non-polarized condition to ensure maximum efficacy with minimal active HOCl generation using an e-bandage. Following HOCl dissipation, H_2_O_2_ can potentially be used for wound healing while maintaining bactericidal activity. Alternatively, H_2_O_2_-producing e-bandages can be employed strategically to eliminate residual HOCl which may damage the host tissue (42). It is important to note that treatment duration cannot be directly extrapolated from *in vitro* to *in vivo* conditions. Several factors, such as bacterial cells embedded within host tissue, biocide consumption by both host tissue and bacteria, wound exudate, and more, may influence optimal treatment times *in vivo* (54–56). Therefore, ideally, this *in vitro* work should be extended to animal models to determine *in vivo* activity. Finally, intermittent treatment using electrochemically-generated HOCl shows excellent biofilm control. This approach has the potential to be utilized for assessing synergistic effects when combining the e-bandage with other drugs/biocides (57). Another potential application of intermittent treatment is that it provides a period of inactivity, during which a potentiostat, used to power the e-bandage, can be attended to for maintenance tasks such as battery replacement and troubleshooting. This intermittent treatment approach should be fine-tuned and evaluated *in vivo*; a deep learning model could be employed to reduce the number of experiments in future design iterations.

## CONCLUSIONS

In this study, intermittent treatment with an e-bandage that generates HOCl and H_2_O_2_ showed promise for controlling MRSA biofilms. The extent of biocidal activity was correlated with HOCl treatment duration. Immediate electrochemical generation of H_2_O_2_ after HOCl resulted in a detrimental effect to biocidal activity due to interaction between these two oxidizing agents. Furthermore, the HOCl-generating e-bandage polarized for just 3 hours retained activity up to at least 7.5 hours after cessation of device polarization, and retained efficacy in several intermittent delivery strategies.

## ACKNOWLEDGEMENTS

Research reported in this publication was supported by the National Institute of Allergy and Infectious Diseases of the National Institute of Health under award number R01AI091594. The author also acknowledges the support from the NIGMS Biotechnology Training Program (T32 GM 8336). The content of this article is solely the responsibility of the authors and does not represent the official views of the National Institute of Health. Fig. 1 was created with BioRender.com.

## Data availability statement

Data are available from the corresponding author upon request.

