## Supplementary material for "Evaluation of Treatment of Methicillin-Resistant *Staphylococcus aureus* Biofilms with Intermittent Electrochemically-Generated H_2_O_2_ or HOCl": SI files

Haluk Beyenal, Ph.D. (Professor)

Associate Dean for Research and Graduate Studies

School of Chemical Engineering and Bioengineering

Voiland College of Engineering and Architecture

Washington State University

.

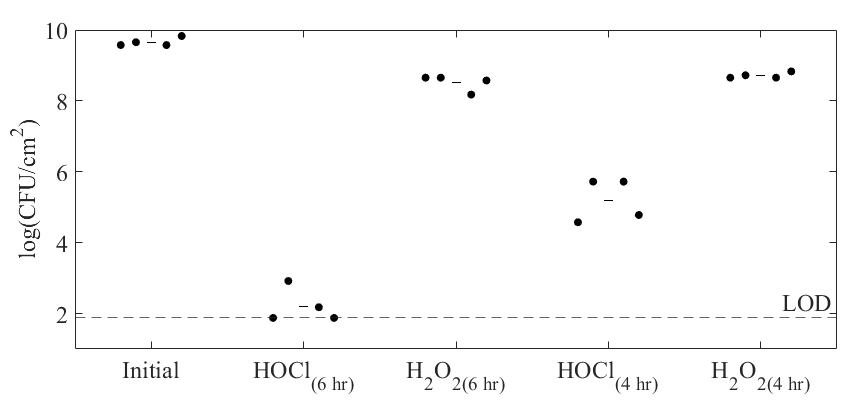

**Figure S1.** **Comparison of results of 4- and 6-hour treatments with HOCl- or H_2_O_2_-generating e-bandages**. Biofilm cell counts are shown as log CFU/cm^2^. Data is represented as individual data points (circles) and means (lines) of at least four independent biological replicates. Six-hour treatment with HOCl e-bandage resulted in 7.45 ± 0.52 log CFU/cm^2^ reduction, which falls within the limit of detection (LOD). Based on this finding, 6 hours of treatment was selected to evaluate the effects of intermittent treatment using HOCl- or H_2_O_2_-generating e-bandages.
